# Characterization of gelling agents in callus inducing media: Physical properties and their effect on callus development

**DOI:** 10.1101/2024.02.14.580317

**Authors:** Noy Sadot Muzika, Tamir Kamai, Leor Eshed Williams, Maya Kleiman

## Abstract

In plant tissue culture, callus formation serves as a crucial mechanism for regenerating entire plants, enabling the differentiation of diverse tissues. Researchers have extensively studied the influence of media composition, particularly plant growth regulators, on callus behavior. However, the impact of the physical properties of the media, a well-established factor in mammalian cell studies, has received limited attention in the context of plant tissue culture. Previous research has highlighted the significance of gelling agents in affecting callus growth and differentiation, with agar, Phytagel, and Gelrite being the most used options. Despite their widespread use, a comprehensive comparison of their physical properties and their subsequent effects on callus behavior remains lacking. Our study seeks to bridge this gap by providing a thorough analysis of the physical properties of these gelling agents and their influence on callus induction and differentiation, offering valuable insights for researchers seeking to optimize plant tissue culture media for diverse applications.

**Highlight:** Studying the impact of gelling agents: Comprehensive analysis reveals how the physical properties of the growing media alters plant callus phenotype and differentiation in tissue culture.

## Introduction

Plant callus is a mass of proliferating cells that are not organized as a tissue and exhibit characteristics of parenchymatous cells^1^. In nature, callus formation is triggered by injury or damage which can be caused by factors such as wind or herbivory^2^. This mass of cells, predominantly composed of pluripotent cells, acts as a protective mechanism, potentially leading to tissue regeneration and the healing of wounded sites^3^. In tissue culture systems, callus is induced by culturing an explant on a solidified medium^4^ supplemented with growth regulators^5^, typically auxins and cytokinins^6^, to promote cell division. The balance between auxins and cytokinins in the medium will determine whether the callus continues to proliferate or regenerate. Higher auxin concentrations tend to promote root formation, while cytokinin promotes shoot regeneration^7^.

Calli differ in their phenotype, texture, gene expression patterns, rates of cell division, degrees of differentiation, and most notably, in their regeneration potential^8^. Understanding and controlling the type of callus that forms is crucial in tissue culture for successful plant regeneration and other biotechnological applications. The type of callus that forms can depend on a variety of factors such as the plant species, the type of explant used^9^ (the tissue from which the callus originates), and the composition of the growth medium^10,11^, which includes the specific types of auxin and cytokinin hormones^12^ as well as their ratio.

In addition to the content of the growth medium, the gelling agent could potentially have major effects on the callus type, as it determines the medium’s physical properties based on the specific arrangement that is formed during the gelling process. This is achieved through the creation of a three-dimensional network of gelling agent’s polymer chains throughout the liquid, resulting in a structure that is more solid than liquid^13^. Consequently, the media possesses a certain rigidity while still retaining the moisture content of the original liquid.

In plant tissue culture, there are three commonly used gelling agents. Agar, is the most widely used gelling agent, owing to its availability, low cost, and good gel strength^14^. It is extracted from the cell walls of red algae, primarily Gelidium (Rhodophyta)^15,16^, and consists mainly of two polysaccharides: agarose, which confers gelling properties, and agaropectin, which provides thickening properties^17^. It is a semi-translucent, hydrophilic colloid that forms strong gels upon heating due to the formation of coiled helices, which effectively retain water molecules^18^. Phytagel^TM^ and Gelrite^TM^ are both derived from gellan gum^19,20^ an anionic linear polysaccharide produced by the bacterium *Pseudomonas elodea*^21^. The gellan gum is composed of glucose, glucuronic acid, and rhamnose. Both gelling agents form a high-strength gel at relatively low concentrations^22^. They also produce colourless media, which is transparent and clear enabling better detection of microbial contamination and facilitating observation of the plant^23,24^.

Though the hydrogels formed by those gelling agents demonstrate distinct properties, a comprehensive characterization and investigation of their effects on responses in plant tissue culture are limited. The frequency of regeneration of Poa pratensis L. in media solidified with Gelrite was twice as high compared to agar^25^. Similarly, the occurrence of callus induction in two rice cultivars was significantly higher in media with Phytagel and Gelrite^26^. However, the specific properties of the media leading to these results have not been addressed in this work. Here we study the gelling agent’s physical and mechanical properties and their effect on Arabidopsis callus behaviour. As the hydrogels formed by the different gelling agents at different concentration produce gels with varying degrees of porosity, ability to hold water and toughness, we evaluated these parameters and correlated them to callus development. We hypothesized that porosity could affect the mobility of water, nutrients, and hormones, thus determining the access of cells cultured on the medium to these essentials. We also hypothesized that the mechanical properties of the hydrogel can change the morphogenetic responses of cells leading to distinct fate, a known phenomenon in mammalian cells^27^. We found that while the hydrogels formed by different gelling agents at different concentrations indeed vary in those properties, only the mechanical properties of the hydrogel correlated with callus type and development.

## Materials and methods

### Callus Inducing Media (CIM) preparation

Callus inducing media was prepared as follows: 3.2gr of Murashige and Skoog B5 (MS) media mix (G5893, Sigma-Merck), 0.5gr of MES salt (M8250, Sigma-Merck) and 2% (20gr) sucrose were added to 900 ml distilled water and stirred thoroughly. pH was adjusted to 5.75 using KOH 1M. Gelling agents (Phytagel (P8169, Sigma-Merck), Gelrite (G1101.0500 Duchefa) or agar (9002-18-0, Merck)) were added while stirring at desired concentration and water was added to a total of 1L. The mixture was autoclaved and cooled to 60°C. At this point kinetin (K3253, Sigma-Merck) 0.1mg/L and 2, 4-D (D7299, Sigma-Merck) 0.5mg/L were added. Media was then poured to a petri dish in a biological hood and the dish remained there until completely cooled.

### Growing conditions and Plant material

In this study, the plant materials used were Arabidopsis accession Columbia-0 (Col-0) and a transgenic plant carrying the *DR5::GFP* construct. Seeds were surface sterilized and sown on plates containing Murashige and Skoog medium^3^ (Sigma-Merck, G5893). The plates were placed for 3 days at 4°C and then transferred to a long day (16h light/8h dark) growth room at temperatures of at 20°C. After one month, leaves number 3-4 were cut at the petiole and placed, abaxial side facing down, on plates with CIM with different gelling agents. The plates were placed in the growth chamber at 20°C until callus growth was observed (about two weeks).

### Callus induction and characterization

WT *Arabidopsis* thaliana leaves number 3-4 were placed on CIM plates. There were 3 plates for each condition (specific gelling agent concentration), each plate had 18 leaves, giving 54 leaves for each condition in total. Successful calli initiations (80%-100%) were characterized at 21- and 28-days post culturing using stereomicroscope (Nikon SMZ25, X5 magnification) and viewed using NIS-elements program (Nikon). Compact or friable phenotype was given to each callus based on appearance. In addition, root hairs were given a score between 0 (no root hairs) to 5 (many root hairs) to quantify root hair growth (see supplementary fig. 3).

### Density measurements

Cubes (1×1×0.5cm) were cut from a petri dish with 25ml solid media using a cuvette (Sarstedt No.67.755). The cubes were then carefully weighted.

### Water loss and gain

Standard petri dishes, 9cm diameter, were filled with 25ml of media. The dishes were weighted, and recorded (M_w_). Dishes were placed at 37°C for 24 hours to dry and weighted again (M_d_). Percentages of weight loss were calculated as the difference between the two weights divided by the initial weight as follows: 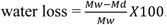. The plates were then submerged in distilled water for an additional 24 hours. Excess water was removed from the plates by carefully tilting, plates were then weighted again (M_rw_). The percentage of weight gain was calculated as follows: 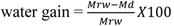. Three repeats for each media (0.8%, 1.2% agar, 0.3%, 0.5% Phytagel, 0.25%, 0.37% Gelrite) were performed.

### Capillary movement

10cm x 1cm strips of Whatman paper were marked with ten lines at 1cm intervals. The edge of the strip was folded and placed on top of the media. The remainder of the strip was placed perpendicular to the media with water visibly moving along the strip. After 10 minutes, the height of the water line was measured. Three replicates were performed for each media.

### Surface humidity

Round Whatman paper, 9cm diameter (GE Healthcare) was weighed and placed on top of the media in a 9cm petri dish (25ml of media). The paper was removed after a minute, weighted, and then placed back on top of the media, repeating this process five times in total.

### Compression tests

Petri dishes of 3cm diameter with 8ml of media were left open in a biological hood until fully solidified. The media was then placed in a texture analyzer (Stable Micro Systems, TA. XT *PlusC*). Bloom test for gelatins was set - a P/20-cylinder probe penetrated 4mm into the gel. Force applied was recorded continually. Five samples were tested for each of the six media types.

### Cryo-SEM

Cylinder of media (2.5mm) was cut from a petri dish and frozen using liquid nitrogen. The frozen samples were sublimated at −85°C for 10 minutes and transferred to a SEM chamber (Joel JSM-7800F scanning electron microscope), where it was carefully cleaved using a scalpel to expose the inner structure. The low temperature (−80°C) of the sample was maintained throughout the imaging process to prevent damage or alteration of the sample.

### Callus induction as response to hormones diffusion

B5 media (35ml) with no hormones was poured into square, gridded petri dishes (10X10cm) and solidified in a biological hood. Using the wide side of a 200µl pipette tip, the media was punctured in the center of the plate. 25µl of kinetin (1mg/ml, Sigma-Merck) and 25µl of 2,4-D (5mg/ml, Sigma-Merck) were mixed and placed in the hole created. *Arabidopsis thaliana* Colombia *DR5: GFP* leaves number 3-4 were cut at the petiole and placed 1cm, 2cm and 4cm away from the center of the plate (fig. 4D). The plates were covered and placed at a growth chamber at 20°C for 3 days to observe hormone disperse. Observations were done using stereomicroscope (Nikon SMZ25, X10 magnification) and viewed using NIS-elements program (Nikon). GFP (395 nm emission) was found in cells containing the hormone auxin, indicating that the hormone has traveled through the media, reaching the leaves, and initiating callus.

### Color disperses

Petri dishes of 15cm diameter with 50ml of media were left to dry in a biological hood. Using the wide side of a sterile 200µl pipet tip, a hole was punctured in the center of the plate. Blue food coloring (50µl) (Maimon’s Brilliant blue FCF, E133) was carefully dripped into the hole. Images of the plates were taken at 24 hours intervals for 96 hours. The camera used is Nikon D3400 N1510, placed 25cm vertically from the plates. The room was lit using fluorescent lighting. Three plates were used for each of the six media types.

### Image analysis

Color disperse images were analyzed with the Image Processing Toolbox (IPT) in Matlab (R2020b) in three steps:

#### 1. Determining the center of the petri dish

The center of the petri dish was determined by relying on the grid of the dish cap visible in the image, with the assumption that the center junction of the gridlines is aligned with the center of the dish. The raw RGB image, obtained from the camera, was transformed into a grayscale image using the *rgb2gray* function (IPT). Subsequently, the *detectLines* function (Matlab exchange) was used to detect the lines in the image, while lines more than 5° from the vertical and horizontal directions were filtered out. Finally, the center of the petri dish was identified as the center point of the intersections of these lines.

##### Correlating between the physical spatial scale and the pixels’ size and location

The correlation between the pixel size in the image and the corresponding physical dimension was established separately for each image, accounting for potential variations in the distance between the camera and the dish. A 6-cm-long rectangle positioned adjacent to the petri dish was utilized as the scale in all the images for this purpose. Figure 4A presents an example of an image with both the center and the rectangle distinctly marked. This process involved the utilization of the raw RGB image, which underwent linearization and transformation into grayscale using the *rgb2lin* and *rgb2gray* functions (IPT), respectively. Subsequently, the edges in the grayscale image were detected using the *edge* function (IPT) with the ‘cany’ option, and regions proximate to these edges were filled using the *imfill* function (IPT), with small regions being excluded from this group through the *bwareaopen* function (IPT). The regions were then labeled using the *bwlabel* function (IPT), and their outer dimensions were measured using the *regionprops* function with the ‘BoundingBox’ option (IPT).

Using the known expected height of the rectangle used for scale, that region was detected by searching for a range of the expected scale. Finally, the pixel-to-dimension scale of the image was determined by dividing the number of pixels of the height of the rectangle by 6 cm.

##### Analyzing the colored image for intensity

The intensity of the brilliant blue color within the specified region of the petri dish (refer to Fig 4A) was analyzed. The ‘saturation’ level of the image was determined by converting the original RGB color scheme to HSV using the rgb2hsv function (IPT). Information was then derived from the second channel (HSV: hue, saturation, value). A radial distribution was assumed, with the origin centered in the petri dish, leading to the transformation of the coordinate system from Cartesian to radial (i.e., from x-y to r-angle). Pixels with identical radial distances (varying angles) were subsequently grouped. For each radial distance, multiple ‘saturation’ values were observed, from which the mean and standard deviation were ascertained (see Fig 4B).

##### Estimating the diffusion coefficient

Considering a finite amount of color applied during a short time relative to the duration of the experiment, and a substrate thickness relatively small to its lateral dimension (radius of the petri dish), the application of the color to the petri dish was approximate as an instantaneous line source (i.e., homogeneous in the vertical center direction of the petri dish).

The analogy is employed between thermal diffusivity, denoted as κ (m2 s⁻¹), and the diffusion of color within the medium. When a finite amount of heat, represented as Q (J m⁻¹), is instantaneously introduced from a linear source positioned within an unbounded medium, the alteration in temperature (ΔT) at a distance r (m) from the source, at time t (s) ^28^:

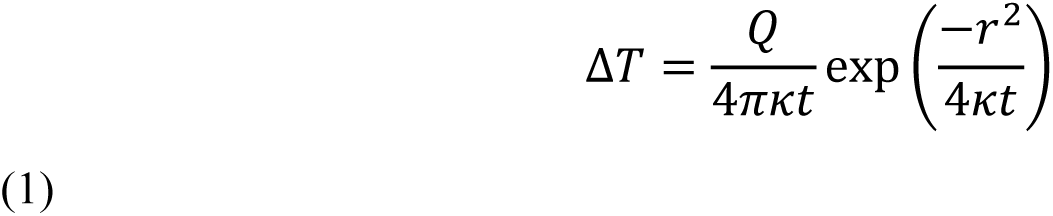

The saturation level extracted from the HSV image is utilized as a measure for color spreading, with the inherent assumption that it is linearly correlated with color intensity and can be quantified. While the saturation level is analogous to ΔT of the model (Eq. 1), a direct quantifiable measure for estimating *Q* from the applied color quantity is not available. Therefore, both parameters, κ and *Q*, were optimized during the fitting of the model’s ΔT to the saturation level observed in the data. However, while a unique κ value was estimated for each image, a shared *Q* value that best fits the data from the transient series of images captured at 24, 48, 72, and 96 hours following the color application was sought.

## Results and discussion

### Callus phenotypes

In mammalian cell culture systems, the structure and physical properties of the media can significantly affect various aspects of cell growth, behavior, and productivity, such as cell proliferation, morphology, differentiation, viability, adhesion, and more^27,29,30^. The composition of the gelling agent on which plant cells are cultured has also shown effect on Tobacco explant shoot regeneration^31^ and it was found that varying gelling agent components increased regeneration. To test how different media affect callus development, we cultured *Arabidopsis thaliana* leaves, on callus-inducing media (CIM) with different gelling agents at their standard concentrations in grams per liter and at concentrations 1.5 times higher. We then categorized the induced calli based on their surface phenotype as either “friable” or “compact”^32^. Friable callus, represented by a 21-day-old 1.2% agar callus (Fig. 1A), is loosely organized, and has a soft and crumbly texture. The cells are typically less packed and can break apart, probably due to weaker cell wall adherence. Compact callus, represented by a 21-day-old 0.25% Gelrite callus, (Fig. 1A), is more structured, with a firmer and more unified texture having a smooth surface. Compact callus is usually characterized by smaller cells that are closely packed together^33^. We characterized 54 calli on day 21 and 28 for each type of medium (Fig. 1A) and observed significant differences in their friable/compact appearance phenotypes. (Fig. 1B). Only 10% of the calli cultured on 1.2% agar exhibited compact phenotype whereas those cultured on 0.25% Gelrite demonstrated approximately 60% incidence of the compact phenotype. In general, calli cultured on media solidified with a gellan gum-derived gelling agent displayed a higher percentage of compact calli than those cultured on media solidified with agar. Differences in callus behavior, leading to either compact or friable phenotypes, were shown to be affected by variations in nutrients and growth regulators in the media^34^. One study showed that in coffee, a higher concentration of Phytagel in the culture medium led to a greater percentage of compact calli and subsequently resulted in a larger number of somatic embryos (71.3) compared to the friable ones (29.2)^35^. Similarly, but in a wider manner, here, the gelling agent is the sole factor affecting the texture of the calli. The success of various callus applications, including cell suspension cultures, somatic embryogenesis, and regeneration, depends significantly on the callus texture. For example, friable calli are favored for cell suspension cultures due to their loosely attached cells, which make them easy to break apart and their rapid cell division^32,36,37^. Therefore, the ability to influence and guide the callus towards a specific texture offers a significant advantage.

**Figure 1.**
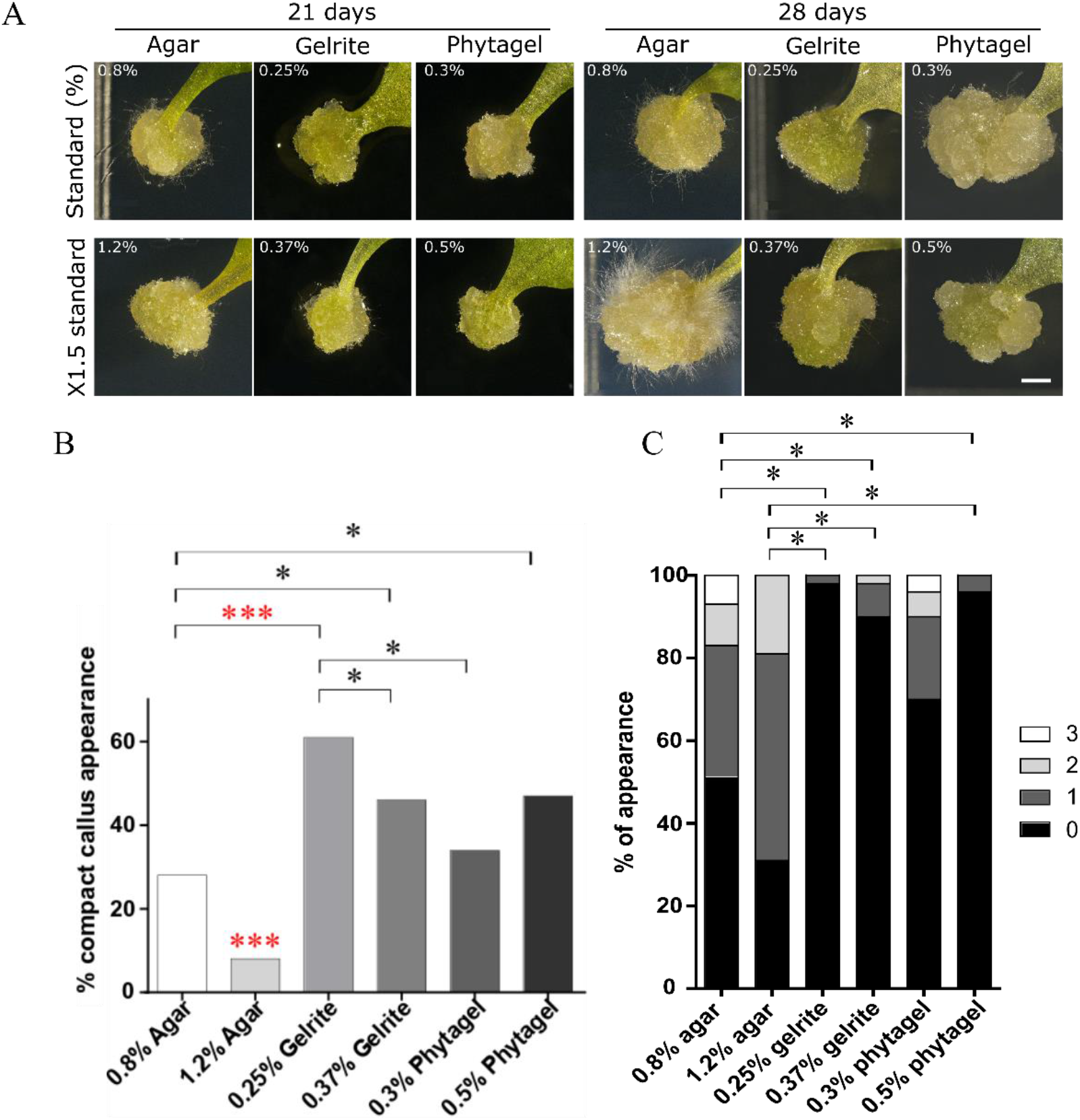
Callus phenotype on media solidified with different gelling agents at different percentages. **A)** Images of callus developed on leaf explants after culturing for 21 and 28 days on callus-inducing media (CIM) containing different gelling agents. Upper panel: Standard concentration expressed as a percentage for each gelling agent. Lower panel 150% of standard. Callus grown on 1.2% agar CIM exhibited the highest number of root hairs. Scale bar = 1 mm. **B)** Percentage of calli showing compact phenotype after 21 days of culturing. Calli were categorized as “compact” or “friable” based on their surface texture. Calli grown on 1.2% agar CIM showed significantly lower numbers of compact calli compared to all other calli (p < 0.0001). (n=34-51) per media. **C)** Percentage of calli showing root hair differentiation after 21 days of culturing. Calli were scored on a scale from 0 (no root hairs) to 3 (many root hairs). Calli on 0.25% Gelrite and 0.5% Phytagel exhibited the lowest percentage of root hair development. All statistical analysis was done using odds ratio and Steel Dwass comparison for all pairs in JMP. (n=34-54) per media.

Root hair appearance on the callus surface indicates cell differentiation. To evaluate whether the type of media affects cell differentiation, we scored each callus on a scale from 0 (no root hairs) to 3 (abundant root hairs). Our analysis revealed significant differences in root hair development among calli cultured with different gelling agents (Fig. 1C). Remarkably, calli grown on 1.2% agar displayed the highest root hair differentiation, while those cultured on 0.25% Gelrite showed the lowest, with very few calli exhibiting low root hair differentiation. In Pearson’s correlation analysis, examining the average root hair severity and the percentage of compact calli, we obtained a p-value of 0.04, indicating a strong association between compact callus appearance and fewer root hairs, and conversely, between friable callus appearance and more root hairs. Cell differentiation can impair the capacity to regenerate as cells lose their potency and the capacity to divide.

Our results demonstrate that altering the gelling agent in the medium, without changing nutrient or hormone levels, directly impacts callus behavior and differentiation. The scope of gelling agents encompasses various factors that can affect cell differentiation and callus behavior. The callus is a three-dimensional structure that forms on a flat surface of the media. Therefore, apart from the physical interaction of a small fraction of the callus with the media that can influence cell behavior, other features of the media, such as water retention, nutrient and hormone movement, gas exchange, and media toughness, might influence callus maintenance and behavior.

We conducted a series of experiments to characterize the physical properties of the media solidified by the different gelling agents. Our goal was to isolate the properties responsible for changes in callus behavior and differentiation.

### Analyzing Water-Related Characteristics of Media with Various Gelling Agents

Water is a critical factor in plant growth and development. In a tissue culture system, the only source of water comes from the hydrogel, which highlights the importance of the hydrogel’s water in maintaining cell turgor pressure for cell health, rigidity, proliferation, and differentiation, as well as facilitating the transport of nutrients and waste removal, essential for sustained cellular activity and growth over time^38^. In addition, water in the medium helps maintain a stable pH, essential for enzymatic activities and overall cellular function. Therefore, the media capacity to hold and release water can affect plant growth *in vitro*. To understand how each gelling agent influences the water-related properties of the media, which may subsequently alter callus behavior, we examined key aspects such as the density, water retention, and surface moisture in media solidified with various gelling agents. This approach aims to reveal the specific ways in which these agents interact with water within the medium and how these interactions might affect callus development.

First, we measured the density of the hydrogel, by weighing three identical 1 cm x 1 cm x 0.5 cm cubes from each type of hydrogel (Fig. 2A). Cubes from 0.25% Gelrite exhibited the lowest average weight, at about 0.3 grams per cube. In comparison, cubes made from agar-based media weighed more, each exceeding 0.4 grams. Since all cube volumes were identical, the differences in cube weight graphed in Fig. 2A are proportional to the differences in media density. The density of the hydrogel is proportional to its water content; a higher density generally indicates a more compact structure with less capacity for water absorption and retention, while a lower density typically allows for greater water uptake and swelling^39^. The most distinct differences in callus phenotype were observed with 1.2% agar and 0.25% Gelrite gelling agents, with agar-grown calli being friable and Gelrite-grown calli remaining compact. The same pattern was observed for root hair differentiation, where calli from 1.2% agar displayed the most root hairs overall. We hypothesize that calli grown on 0.25% Gelrite, having lower density, may have higher access to water, nutrients, and hormones, thus displaying compact phenotype. Conversely, denser media (1.2% agar) could require active nutrient and hormone extraction, potentially limiting availability and affecting cell pluripotency.

**Figure 2.**
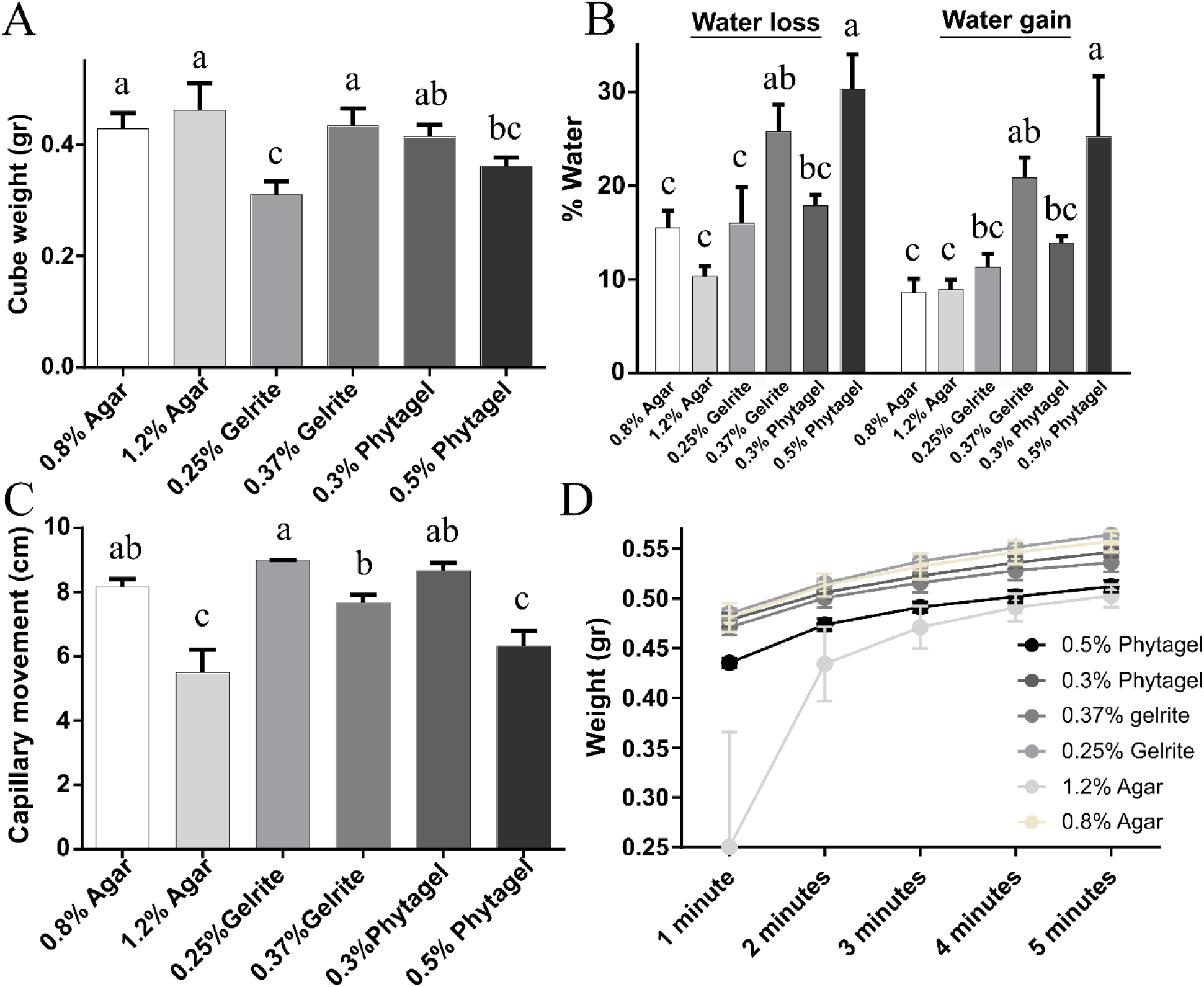
Differences in water content and surface humidity of callus inducing media (CIMs) solidified with distinct gelling agents at different concentrations. **A)** Media density. Cubes of 1×1×0.5 cm were cut out of the media and weighted to determine the average weight (n=3). **B)** Media water retention and post-drying re-absorption. Plates with media were weighed (Mw) and then placed at 37°C for 24 hours to dry, weighed (Md), and then submerged in distilled water for an additional 24 hours. Following the removal of excess water, the plates were reweighed (Mrw). Water loss and water gain were calculated as percentages [(Mw-Md)/Mw X100] and [(Mrw-Md)/Mrw X100], respectively (n=3). **C)** Capillary movement on CIM surface. A strip of Whatman paper was placed on the surface of the CIMs for 10 minutes, and the distance of water traveling upstream was measured (n=3). **D)** Media surface wetness. A standard 9cm Whatman paper disk was weighed and placed on the surface of the media. The paper was weighed every minute for 5 minutes, and the weight was normalized as [(Ws-Wt)/Ws], where Ws is the start weight and Wt is the weight at a given time point (n=3). The results show that 0.25% Gelrite had the highest water content and surface wetness, while 1.2% agar had the lowest in both cases. Statistical analyses were done using Tukey-Kramer HSD.

One of the features that gelling agent can determine is water retention, which influence the rate of water loss. Hence, evaluating the water loss can indicate water retention capacity of the hydrogel. A high rate of water loss might suggest that the hydrogel has a lower capacity to retain water, which could be a result of its composition, cross-linking density, or porosity^39^. Following standard protocols of hydrogel characterization^39^, we measured water loss after exposure to 37°C for 24 hours and water regain of the different media. Media with 0.5% Phytagel exhibited the highest water loss, losing nearly 30% of its original weight and recovering most of it upon rehydration. Media with 0.37% Gelrite exhibited a similar pattern, losing about 25% of its original weight and gaining back about 80% of it after rehydration. In contrast, 1.2% agar media, lost the least amount of water when dried, approximately 15% of its original weight, and regained roughly 66% of it (Fig. 2B). Water loss and gain exhibit similar trends across the media, showing for the reproducibility of our results. Water retention and water holding has been suggested as an important factor in plant growth and hence several studies in recent years suggested the addition of hydrogel to soil to increase water retention^40^. Changes in the polymer chemistry, particle size and network structure were altered to vary the water retention ability. While we did not investigate the reason behind the change in water retention in our media, it should be noted that we found no correlation between water-holding capacity and callus phenotypes, indicating this property has minimal impact on callus behavior.

While water retention showed no correlation with callus behavior, we suspected that water availability at the point of contact between the plant and the media, which is the plant surface, may influence callus phenotype. It is widely recognized that plant tissue must overcome the capillary force retaining water within the media for water uptake^41^. To test whether different hydrogels vary in their capillary force, we measured the upward capillary movement along a Whatman paper strip within 10 minutes (Fig. 2C, supplementary Fig. 2A). Our results showed that 0.25% Gelrite exhibited the greatest capillary movement, with water reaching the end of the strip after 10 minutes. Conversely, 1.2% agar and 0.5% Phytagel exhibited reduced capillary movement, suggesting that a stronger force is required to draw water over their surfaces. Furthermore, we measured the surface wetness of each medium by placing a standard 9cm Whatman paper disk on the surface and recording the weight change over time as it absorbed water (Fig. 2D, supplementary Fig. 1B). We observed that papers on 0.25% Gelrite gained the most weight and did so the fastest, while papers on 1.2% agar and 0.5% Phytagel gained the least weight over time. Although no significant weight gain was observed, the hydrogel containing 1.2% agar exhibited both the slowest rate of increase and the lowest overall weight gain. These results suggest that 0.25% Gelrite hydrogel has the highest initial water availability and surface wetness, while 1.2% agar has the lowest. This suggests that water availability upon direct contact may play a role in maintaining callus identity.

### Structural analysis – compressibility and porosity

Some plants grown in tissue culture systems respond to media elasticity and compressability^42^. To characterize those properties in the media we used, we conducted both compression and oscillatory shear stress tests. In media compressibility, we measured the force applied to compress the hydrogel by 4 mm (Bloom’s test for gels, Fig. 3A). As the applied force is higher, the media is less compressible. The media prepared with 1.2% agar was the least compressible, requiring a maximum force of about 800gr to be compressed by 4mm. The 0.25% Gelrite media resisted the least with a value of about 50gr force needed for penetration. A higher concentration of the gelling agent required a force about 2-3-fold greater to compress by 4 mm (ration in: Agar – 3.29, Gelrite – 2.97, Phytagel – 2.33). To characterize the viscoelastic behavior of the hydrogel we tested oscillatory shear stress (Fig. 3B). In this test, we apply increasing values of shear stress to test at which point the material changes from a predominantly elastic material (that bounces back to its original form upon deformation) to a predominantly viscous material (that flows and deforms upon applied pressure). We measured the cross point between G’ (elastic modulus) and G’’ (viscous modulus) values also known as the gelling point - the pressure under which the material transitions from elastic to viscous. This is, in fact, a measurement for the strength of the cross-linked network while deforming the media. In agreement with earlier work, media containing 1.2% agar was the most difficult to break down^43^ suggesting a strong cross-linked network. The 0.25% Gelrite was the easiest to break, showing the weakest total cross-link bonds. The oscillatory shear stress results align perfectly with the compression results. This is due to the homogenous nature of our materials that do not have a preferred axis with stronger cross-link bonds^44^. These results also correlate with callus differentiation tested; on less compressible media, calli tended to differentiate faster, while on weak cross-linked network media, calli tended to keep their callus identity. Testing the correlation between the percentage of compact callus appearance and the compression value of the hydrogels resulted in a Pearson’s correlation coefficient (r) value of 0.7 (p-value = 0.03), indicating that there is a statistically significant relationship between these two variables (Figure 3C). Compact calli developed on less compressible media, while friable calli on tougher media. This response to the media material properties is well known and has been established in mammalian cells^45^, however, this is the first time we see such a correlation in plant cells. Next we employed cryo-scanning electron microscope (cryo-SEM)^46^ to visualize and make comparisons regarding the structure of various hydrogels (Fig. 3C). We observed that all hydrogels have oval pore-like networks that are relatively uniform in size throughout the media. Pore size varies significantly between materials, depending on type and concentration of the gelling agent^47,48^. To quantify these differences, we measured the diameter of 90 pores in our SEM images using the “measure” function in FIJI (Fig. 3D). While the smallest pore size measured was less than 2µm in 1.2% agar media, the largest pores were measured for 0.37% Gelrite media at an average of 6.5µm. Remarkably, we observed different trends among gelling agents as their concentrations increased. While in agar, the pore size decreased dramatically from 1.2% to 0.8%, the pore size increased as Gelrite concentration increased. These results do not align with the mechanical properties of the different materials, suggesting that the porosity is not the underlying cause for the differences in mechanical properties. The relation between hydrogel porosity and mechanical properties is not well established and is dependent on many other factors such as chemistry, structure and composition^49^. The same is true regarding water retention, where porosity plays an important role, but other factors such as cross-link density and polymer chemistry need to be considered, which may explain why we do not see correlation between porosity and water retention in our system. A correlation test showed that media porosity does not correlate with the callus phenotypes that we observed. For example, pore sizes for 0.8% agar media and 0.25% Gelrite are very similar and not significantly different from each other, while calli grown on these different media do have a large difference in root hair occurrence and surface phenotype (See supplementary Fig.3). Additionally, despite 0.37% Gelrite having the largest pores, it did not result in distinct callus behavior compared to 0.25% Gelrite. Since there was no correlation between the porosity and the mechanical properties of the material in our system, we were able to separate the two and find the one (mechanical properties) that has an influence on the cultured callus phenotype. However, the size of the pores can influence the uptake of nutrients and hormones by the plant tissue due to capillary forces that retain the water^49^. Smaller pores may limit nutrient and hormone availability, while larger pores can allow for better nutrient penetration. This leads us to explore pore size in relation to diffusion.

**Figure 3.**
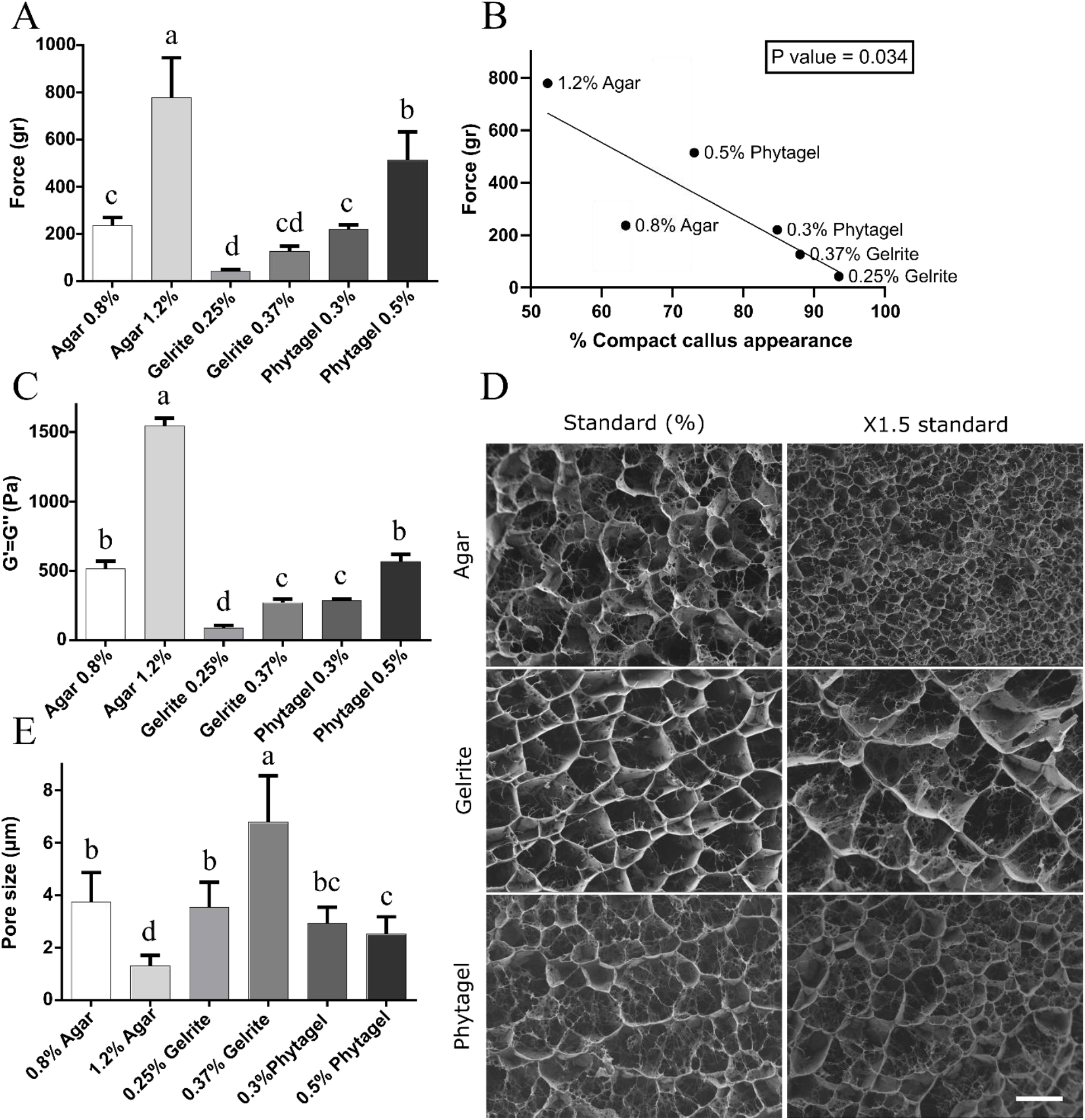
Structural and mechanical analysis of media. **A)** Compression test. The force needed to penetrate 4 mm into the media was measured using a texture analyzer. The 1.2% agar media required the most force, while the 0.25% Gelrite media required the least. **B)** Correlation analysis between the force needed to compress the media, as shown in A, and the compact callus appearance percentage, as shown in Fig. 1B. As media compressibility increases, compact callus appearance decreases (P-value < 0.05). **C)** Shear stress rheology analysis. Oscillatory shear stress was measured up to the point of transformation from elastic to viscous material, where G’ (elastic modulus) equals G’’ (viscous modulus). The 1.2% agar media required the highest stress to reach this point, suggesting a strong, cross-linked network, while the 0.25% Gelrite media showed the lowest values, indicating weaker bonds. **D)** Cryo-scanning electron microscope (SEM) imaging of the media. Media was carefully sliced and extracted vertically from the Petri dishes to preserve structural integrity and ensure consistent orientation. Bar=5 µm. **E**) Pore size analysis based on cryo-SEM imaging. (n=90 pores). Cryo-SEM images were analyzed using FIJI software, “measure” function. All statistical analysis were done in JMP, with comparisons for all pairs using Tukey-Kramer HSD.

### Diffusion analysis

Plant tissues cultured in tissue culture systems rely on the media for minerals and energy in the form of sugars. Since the tissue is in contact only with the surface of the hydrogel, the availability of nutrients, sugars, and hormones—once applied—primarily depends on their ability to diffuse through the hydrogel. To explore potential differences in the diffusion of nutrients and substances in the media, we estimated the relative diffusion coefficient through the spread of food dye over time. We punctured the hydrogel at the center of a 15 cm diameter Petri dish and carefully dispensed 50µl of blue food coloring into the hole. We then allowed the color to spread for over 96 hours, imaging the media every 24 hours (Fig. 4A). Using image analysis, we determined the spread of the food dye over time, by averaging the color saturation in the same distance from the center (black line in Fig. 4B). This average and its standard deviation (grey area in Fig. 4B) were fitted to a radial distribution model (red line in Fig. 4B) to estimate the diffusion coefficient of the blue dye in the media. Consideration of different time points resulted in different diffusion coefficients. All obtained diffusion coefficients were compared to find the terms for a significant difference in the food dye diffusion coefficient in the different media.

**Figure 4.**
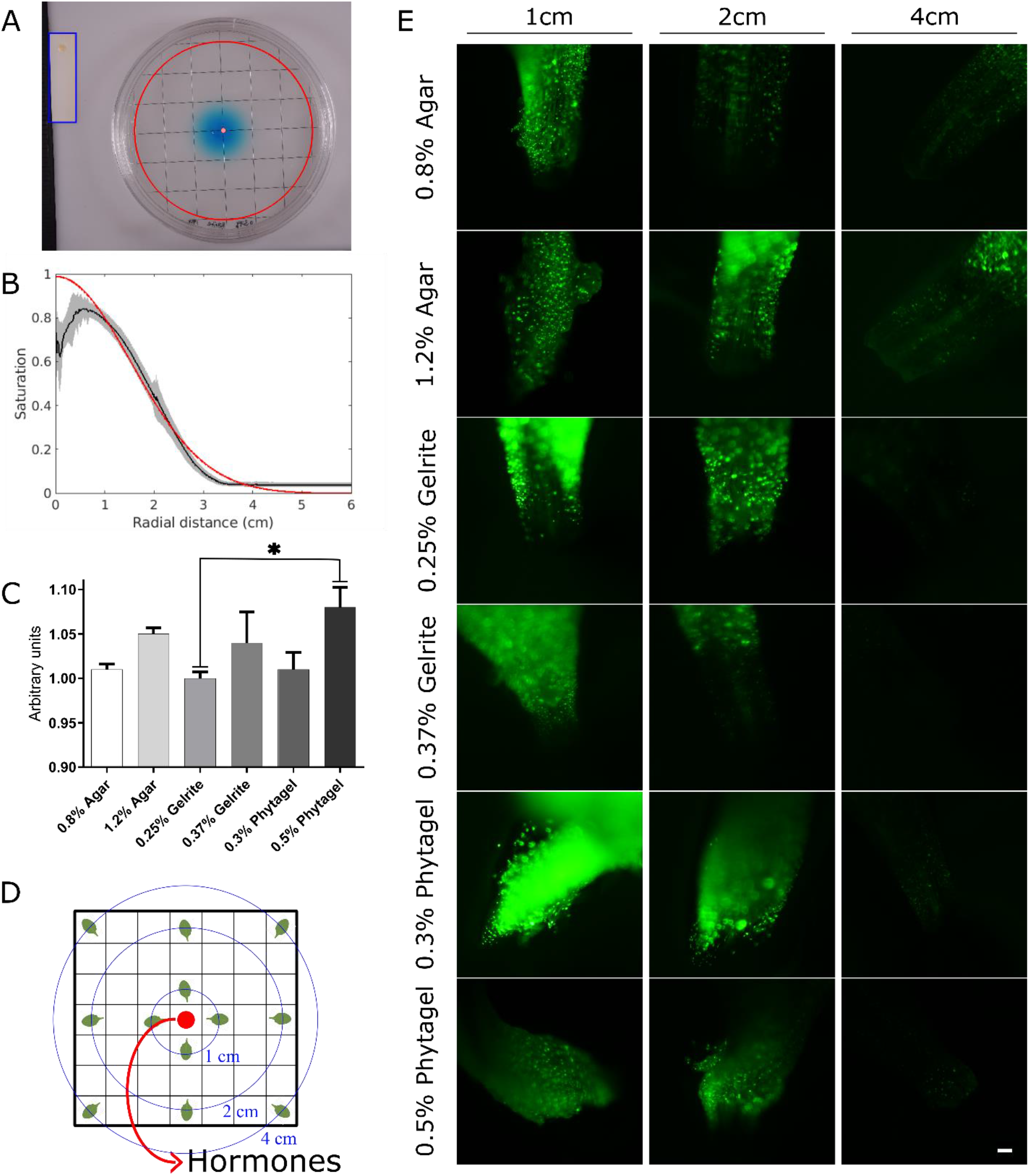
Substances diffusion analysis. A) Blue food coloring disperses on 0.37% Gelrite media at 24 hours. 50µl droplets of blue food coloring were placed in the center (marked by a red dot) of 15cm round CIM plates. The disperse (marked by a red circle) area was observed and measured over 96 hours with 24 hours intervals. Analysis was done for each of the six types of CIM (see supplementary figure 4). B) Radial distribution model and data. The radial diffusion model fit (red line) to the data, indicated by the average saturation at every point in the same distance from the center (black line) and standard deviation (gray area). C) Diffusion coefficient determination. Relative diffusion coefficients as calculated from the model based on the experimental results considering the food dye distribution at time points 24 and 48 hours. D) Schematic visualization of hormones disperse experiment. Leaves are placed on a hormone-depleted media at varying distances from the center (1, 2 and 4 cm). Hormones are then added to the center of the plate. E) Leaves induced calli at different distances from hormonal dose in center plate. 25µl of 2,4-D mixed with 25µl of kinetin were placed in a hole created in the center of a 15cm square gridded petri dish. One month old DR5: GFP *Arabidopsis* leaves were placed at a radius of 1, 2 and 4cm from the hormonal dose. Images were taken at fluorescent light (GFP) 3 days post placement. (n=12 for each distance point, representative images are shown). Bar=1000µm. Statistical analysis was done using JMP, one-way Anova – comparisons for all pairs using Tukey-Kramer HSD.

We received significant differences in the diffusion coefficient of the food dye between the different media only when looking at the time points of 24-48 hours (Fig. 4C). For the analysis, we omitted the initial 24 hours and the time after 48 hour. In the initial 24 hours, the color intensity was relatively high to detect significant differences, while at time beyond 48 hours the dye was too spread out for detecting a distinct line between the dye and the media. We found the largest diffusion coefficient to be in 0.5% Phytagel. However, this was significant only in comparison to 0.25% Gelrite. In the work of Dobránszki et al^42^, diffusion was tested in different blends of Phytagel/agar. They found that methylene blue diffused the fastest in Phytagel, but there are some important distinctions between their work and ours. First, Dobránszki et al^42^ did not calculate the diffusion coefficient, but rather measured the distance to which the dye diffused. This can give contradictory results, as high diffusion coefficient does not necessarily mean longer diffusion distance at a particular time. Second, they used much lower concentration (approximately half) of gelling agent than the concentrations herein, which alters the media properties completely. Third, methylene blue is a salt used as a dye, and not an organic compound like the blue food coloring we used. This is important as the diffusion coefficient is a property of the solute in a specific media and can vary between solutes in the same media. The higher diffusion coefficient found in 0.5% Phytagel and the lower one in 0.25% Gelrite, while significant, do not correlate with the phenotype of the callus, which appears similar. We hence decided to perform the diffusion experiment with the most relevant materials diffusing in the media and influencing callus development - the hormones. To assess the impact of the gelling agents and their concentration on hormone diffusion within the media, we utilized the *DR5::GFP* transgenic plant, a widely recognized tool for visualizing auxin response in live tissues^50^. We placed Arabidopsis leaves on hormone-depleted media at a certain distance from the center of the plate and injected a high dose of auxin and cytokinin (5mg/ml and 1mg/ml, respectively) into the center of the plate (Fig. 4D). After 3 days of incubation, we analyzed the fluorescent signal under a stereomicroscope. All leaves that were placed 1 cm away from the point of injection exhibited a strong signal in all media. At 2 cm, leaves from all media showed a signal, but the one from 0.8% Agar showed the least. At 4 cm, there was mostly no signal, except for a low signal on the leaf placed on 1.2% Agar. This result suggests that hormones diffuse in all hydrogels to the extent that the tissue responds to them. It might be that the hormone concentration we used is too high to discern any differences, or that a quantitative analysis of the fluorescent signal’s intensity is required to detect variations in hormone diffusion.

## Conclusions

Numerous studies have highlighted the significant role of gelling agents in influencing callus, embryo, and organ development in tissue culture systems^25,26,31,51^. In this study, we offer insights into the properties of these gelling agents and their potential contributions to the differential responses observed in tissue cultures.

The most noticeable phenotype among the calli developed on different media was the extensive differentiation of root hairs on the 1.2% agar media. The texture of the calli, whether compact or friable, was also strongly affected, with calli on 1.2% agar showing friable appearance with loosely attached cells and those on Gelrite 0.25% having the most compact appearance. The two media represent hydrogels with the most divergent physical properties across a range of conducted tests. In general, the agar hydrogel is characterized by a high density, as evidenced by its smallest pore size. This feature likely contributes to its low capillary movement, minimal surface humidity, and the force exerted within the medium. These attributes are interconnected, where the small pore size can lead to reduced movement of water and nutrients through capillary action and influence the surface wetness level and the hydrogel mechanical properties. All these factors may reduce the tissue’s ability to effectively uptake water and nutrients, as well as hormones that direct cell proliferation, from the culture medium, resulting in cell differentiation. On the other hand, enhanced water availability in the Gelrite media could facilitate cellular adhesion, potentially leading to the observed compact morphology of the calli. An excess of water can lead to vitrification, a process that potentially disrupts all applications, and therefore should be carefully considered^52^. A future histology analysis can contribute to a deeper understanding of the cellular and tissue-level changes.

Noteworthy is that along with the gelling agent concentration, factors such as pH, ionic strength, and the presence of ions or additives in the culture medium can also significantly affect the properties of hydrogels. For example, both agar and Gelrite media demonstrated softer characteristics when the pH levels were lower, and they exhibited increased hardness at higher pH levels upon introduction into the same medium^23^.

Therefore, understanding the physical properties that gelling agents at different concentrations confer upon the hydrogel can serve as a tool to improve performance.

## Abbreviations

CIM: Callus inducing media

